# Lentivirus-mediated long-term overexpression of specific microRNA for complementary miRNA pairs in mammalian cells

**DOI:** 10.1101/717330

**Authors:** Shupei Qiao, Yufang Zhao, Kai Li, Yulu Sun, Liuke Sun, Xiaolu Hou, Hui Tian, Kangruo Ye, Yongzhan Nie, Weiming Tian

## Abstract

The establishment of a method that would overexpress or suppress of specific microRNA activity is essential for the functional analysis of these molecules and for the development of miRNA therapeutic applications. There already exist excellent ways to inhibit miRNA function in vitro and in vivo by overexpressing miRNA target sequences, which include miRNA ‘decoys’, ‘sponges’, or ‘antagomirs’ that are complementary to an miRNA seed region. Conversely, no methods to induce stable gain-of-function phenotypes for specific miRNAs have, as yet, been reported. Furthermore, the discovery of complementary miRNA pairs raises suspicion regarding the existing methods used for miRNA overexpression. In our study, we will study whether the traditional methods for miRNA overexpression can be used for specific miRNA overexpression while complementary miRNA pairs exist. In addition, we test various miRNA-expression cassettes that were designed to efficiently overexpress specific miRNA through the shRNA lentivirus expression system. We report the optimal conditions that were established for the design of such miRNA-expression cassettes. We finally demonstrate that the miRNA-expression cassettes achieve efficient and long-term overexpression of specific miRNAs. Meanwhile, our results also support the notion that miRNA–miRNA interactions are implicated in potential, mutual regulatory patterns and beyond the seed sequence of miRNA, extensive pairing interactions between a miRNA and its target also lead to target-directed miRNA degradation. Our results indicate that our method offers a simple and efficient means to over-express the specific miRNA with long-term which will be very useful for future studies in miRNA biology, as well as contributed to the development of miRNA-based therapy for clinical applications.

## Introduction

Thousands of non-coding RNAs (ncRNAs) have been observed over the past decade. MicroRNA (miRNA) is a type of ncRNA that is composed of 18–24 nucleotides and is generally highly conserved between different species [1, 2]. There is increasing evidence that miRNAs play important roles in various biological processes, including development, differentiation, and stemness by combining with the 5′UTR, ORF, or 3′UTR of a target gene mRNA to suppress its translation or induce its degradation [3–6]. Moreover, the aberrant expression of miRNA was reportedly associated with human diseases [7–9]. Until now, many studies have contributed to the development of miRNA-based therapy for clinical applications [10, 11]. Many therapeutic miRNAs and small-interfering RNAs (siRNAs) offer important biopharmaceutical properties that have been commercially developed as potential medical treatments [10–13]. The process of analyzing the underlying molecular mechanisms of miRNA regulation is quite extensive. Further, the development of reagents that can strongly suppress and overexpress specific miRNAs has also generated much interest and will be important for basic miRNA research and as a possible therapeutic strategy.

Currently, there are already numerous, well-proven ways to inhibit specific miRNA functions in vitro and in vivo by overexpressing miRNA target sequences, including miRNA ‘decoys’, ‘sponges’, or ‘antagomirs’ that are complementary to an miRNA seed region [14–18]. Among them, the overexpression of efficient RNA decoys through the lentiviral vectors can achieve better effectiveness, which can suppress specific miRNA activity for over 1 month, as compared to the sponges and antagomirs [14]. However, less attention has been paid to specific miRNA overexpression. Several studies have shown that the abnormally low expression of miRNAs usually leads to the upregulation of oncogene, which assumes primary responsibility for the occurrence of cancer [19–22]. At present, the most commonly employed methods to induce miRNA overexpression are by expressing the pre-miRNA via RNA polymerase II or chemically modified double-stranded RNAs that mimic endogenous miRNAs [23–28]. miRNA is typically assumed to be a carrier strand of the mature miRNA; it has demonstrated its effects on gene regulatory networks in both cultured cells and living animals [29]. In addition, an interesting phenomenon was recently discovered, but it remains incompletely understood: Natural or endogenous sense/antisense miRNAs can complementarily bind to each other, such as has-miR-24 and has-miR-3074-5p, has-miR-486-5p and has-miR-486-3p, and so on. The authors of several studies have speculated that the interaction between complementary miRNA pairs implies that there is a complex, potentially mutual regulatory pattern among miRNAs [30, 31]. This discovery has potential impacts on the traditional methods used to overexpress and inhibit miRNAs. In our study, we attempt to expound the shortcomings of the traditional methods employed to study miRNA, and we designed various miRNA-expression cassettes to establish a specific, long-term miRNA overexpression strategy using the shRNA lentivirus system.

## Material and methods

### Lentiviral Vector Construction and Viral Production

Oligonucleotide pairs for miRNA expression cassettes, listed in Supplementary Table S1, were annealed and cloned into the pLKO.1-TRC vector digested with AgeI and EcroRI to generate the pLKO.1-miR lentiviral vector. A human genomic DNA fragment comprising pri-miR-378 and pri-miR-486 was amplified by polymerase chain reaction (PCR) using primers (listed in Table S2). The amplified fragment was cloned into the EcoR I and BamH I restriction enzyme sites of pMD18-T vector (TaKaRa, Kusatsu, Japan) and verified by sequencing. The fragment was excised from the pMD18-T vector and ligated into the pLVSIN CMV Puro destination vector (TaKaRa) resulting in the pLVSIN-miR-378/486 vector. Viral particles were generated by transfecting plated 293T cells with pLKO.1-miR lentiviral vectors or with the pLVSIN-miR-378/486 vector, along with gag/pol and VSV-G vectors at proportions of 9:9:1 using calcium phosphate transfection. After 24 hours, 6 mL of fresh medium was added and the supernatant from the transfected cells was collected at 48 hours and 72 hours. Then, the lentiviral vector was obtained through centrifugation and filtration by a 0.45 µm filter.

### Cell culture and construction of stable cell lines

H460 cells were grown in Roswell Park Memorial Institute (RPMI) 1640 medium, and the SKBR3, A549, HeLa, and 293T cells were grown in Dulbecco’s Modified Eagle’s Medium (DMEM) supplemented with 10% fetal bovine serum (FBS; Thermo Fisher Scientific, Waltham, MA, USA) and 1% antibiotic–antimycotic (Thermo Fisher Scientific). Both were incubated in a humidified atmosphere of 5% CO_2_ at 37°C; the media were changed every other day. The cells were seeded at 5×10^5^ per 10 cm dish and infected with lentivirus, which was packaged by our team within 24 hours; the media were changed after being 24 hours of transduction. The stable cell lines were constructed with the addition of puromycin (5 μg/mL) 48 hours after being infected; for our study, a generation or two of stable cell lines were used in experimental studies due to specific cell behaviors that were generated, such as apoptosis and inhibited cell proliferation, following the introduction of alien genes.

### miRNA–oligonucleotides and transfection

Chemically modified, double-stranded RNAs that mimic endogenous miRNAs were purchased from Shanghai GenePharma Co. Ltd (Shanghai, China) and Thermo Fisher Scientific. Transfection was performed using the siRNA Transfection Reagent (Hoffmann-La Roche, Basel, Switzerland) using miRNA oligonucleotides in OpTi-MEM I (Gibco®; Thermo Fisher Scientific) until they reached a final concentration of 100 nM, as per the manufacturer’s instructions.

### RNA isolation and qRT-PCR

Total RNA was extracted from cell lines using TRIzol reagent (Thermo Fisher Scientific) according to the manufacturer’s instructions; RNA quality was assessed by agarose gel electrophoresis. For the gene expression analysis, a total of 1□μg of RNA for each sample was reverse-transcribed to cDNA using reverse-transcription kits (Promega, Madison, WI, USA), and real-time quantitative reverse transcription (qRT)-PCR was performed on a 7500 real-time PCR System (Applied Biosystems, Foster City, CA, USA) using FastStart Universal SYBR Green Master [Rox] (Hoffmann-La Roche). The primers used in the present study are shown in Table S3. Then, 18S was used as the internal reference for normalization. The relative mRNA expression was analyzed using the 2^−ΔΔCt^ method.

To detect mature miRNA, we performed a stem-loop qRT-PCR assay. miRNAs were reverse transcribed from total RNA using a reverse-transcription system kit (Applied Biosystems) with the miRNA-specific stem loop–reverse transcription (RT) primers, which were designed by our team in accordance with the manufacturer’s instructions. Briefly, 1 µg of total RNA, the specific stem loop-RT primer mix (50 nM for each), 1× RT buffer, 0.25 mM each of dNTPs, 3.33 U/µL of MultiScribe reverse transcriptase, and 0.25 U/µL of RNase inhibitor were intermixed. The 15 µL reactions were incubated for 30 minutes at 16°C, 60 minutes at 42°C, 5 minutes at 85°C, and then held at 4°C. Real-time PCR was performed using the FastStart Universal SYBR Green Master [Rox] (Hoffmann-La Roche) with miRNA forward primer and universal primer (URP). The miRNA-specific stem loop-RT primers and universal primers are listed in Table S4. Real-time PCR was performed on a 7500 real-time PCR System (Applied Biosystems), and the PCR reactions were conducted at 50°C for 2□minutes and 95°C for 10□minutes, followed by 40 cycles at 95°C for 15□seconds and 60°C for 60□seconds using an ABI 7500 fast real-time PCR system. U6 snRNA was used as an internal control, and the relative miRNA expression was analyzed using the 2^−ΔΔCt^ method.

### Cell Proliferation Assay

Cell proliferation capacity was evaluated using the CCK□8 assay, according to the manufacturer’s instructions. The cells from one-or two-generation stable cell lines were seeded onto 96□well plates with the same cell number. CCK□8 (10 μL) was added to each well and the cells were incubated for a further 2 hours at 37 C. The optical density (OD) was measured at 450 nm using an auto□microplate reader (Infinite M200; Tecan, Männedorf, Switzerland).

### Apoptosis Assay and Cell-cycle Assay

After 48□hours, and once selected and enriched by applying puromycin in the culture medium after virus transduction, both the suspension and attached cells were gently collected. The cell density was adjusted to 5×10^6^ cells/mL. Then, 100 μL of cell suspension was incubated with 5 μL AnnexinV/FITC for 10 minutes and then with 5 μL of propidium iodide (PI; BD Biosciences, Franklin Lakes, NJ, USA) for 5 minutes at room temperature in dark. The apoptosis rate was measured by flow cytometry (FCM). For the cell-cycle assay, the cells were trypsinized, harvested, and washed in cold phosphate buffered saline (PBS) once and fixed in 75% ethanol at –20°C for 24□hours. The cells were then centrifuged, washed in cold PBS once, and stained in PI/RNase staining buffer (Tianjin Sungene Biotech Co., Ltd., Tianjin, China) at room temperature for 30 minutes in the dark, at which point they were finally analyzed using a BD FACSCalibur Flow Cytometer (BD Biosciences). For each sample, 10^5^ events were counted.

### Statistical Analysis

The results are presented as the means ± standard deviations. Student’s two-tailed *t*-test was performed to compare the differences between the experimental and control groups. A *P*-value of <0.05(*) was considered significant; a *P*-value <0.01 (**) was considered highly significant.

## Results

### The influence of the discovery of complementary miRNA pairs to the present miRNA overexpression methods

The establishment of an miRNA date-storage platform can provide the miRNA sequence, pre-miRNA secondary structure, miRNA gene loci, and other miRNA annotation information [32, 33]. Many bioinformatics tools have been developed for miRNA biogenesis and to investigate questions within the field of miRNA biology, and many interesting phenomena were subsequently discovered [34, 35]. Remarkably, some miRNA pairs, located in the same or different genomic regions, also showed complementarity, in addition to endogenous sense/antisense miRNAs [30].

### Chemically synthesized miRNA mimics for complementary miRNA pairs overexpression

It is well-known that the miRNA mimics that were used for miRNA overexpression are innovative molecules designed for gene-silencing approaches [36, 37]. Specifically, their design is based on native miRNAs, which usually have mismatches between the 5p and 3p strands. We wonder whether the miRNA mimics that were chemically synthesized are suitable for the overexpression of complementary miRNA pairs. Thus, in our study, we chose the has-miR-486-5p and has-miR-486-3p pair to test our assumption, which is processed by pri-miR-486; these miRNAs were reported to be complementary to each other. As shown in Figure 1, whether transfection with the miR-486-3p mimic or the miR-486-5p mimic in A549 cells all leads to the overexpression of miR-486-3p and miR-486-5p, it is interesting to note that the miR-486-5p expression level is higher than that of miR-486-3p when using the miR-486-3p mimics. To further verify this phenomenon, we performed similar experiments using H460 and HeLa cells; by doing so, we obtained very similar results, except that the expression of miR-486-5p was dramatically lower in HeLa cells than in A549 or H460 cells following transfection of miR-486-3p mimics (Supplementary Figures 1 and 2). Previous studies have shown that miR-486-5p directly targets phosphoinositide-3-kinase (PIK3R1) and protumorigenic ARHGAP5, and that it functions as a potent tumor suppressor of lung cancer both in vitro and in vivo, while overexpression miR-486-3p in cervical cancer cells can inhibited cell growth and metastasis by targeting ECM1 [19, 38, 39]. To study the impact of miRNA on the reported miR-486-regulated gene expression, we detected the expression of *PIK3R1*, *ARHGAP5*, and *ECM1* after transfecting the miR-486-3p or 5p mimics in the A549, H460, and HeLa cells. The results showed that the expression of *ECM1* is significantly downregulated following miR-486-5p mimics transfection in all cell lines, while the miR-486-3p mimics’ transfection also results in the *ECM1* downregulation, but without a remarkable difference, which is inconsistent with the previous report. However, we found that the *PIK3R1* gene can be downregulated by the transfection of the miR-486-3p or miR-486-5p mimics in the A549 or H460 cells, but with no change in ARHGAP5 in the A549 and HeLa that were transfected (Figure 1d; Supplementary Figures 1d and 2d). Based on these results, we may conclude that while the miRNA we studied are complementary to each other, use of chemically produced miRNA–oligonucleotides should be minimized.

**Figure 1.**
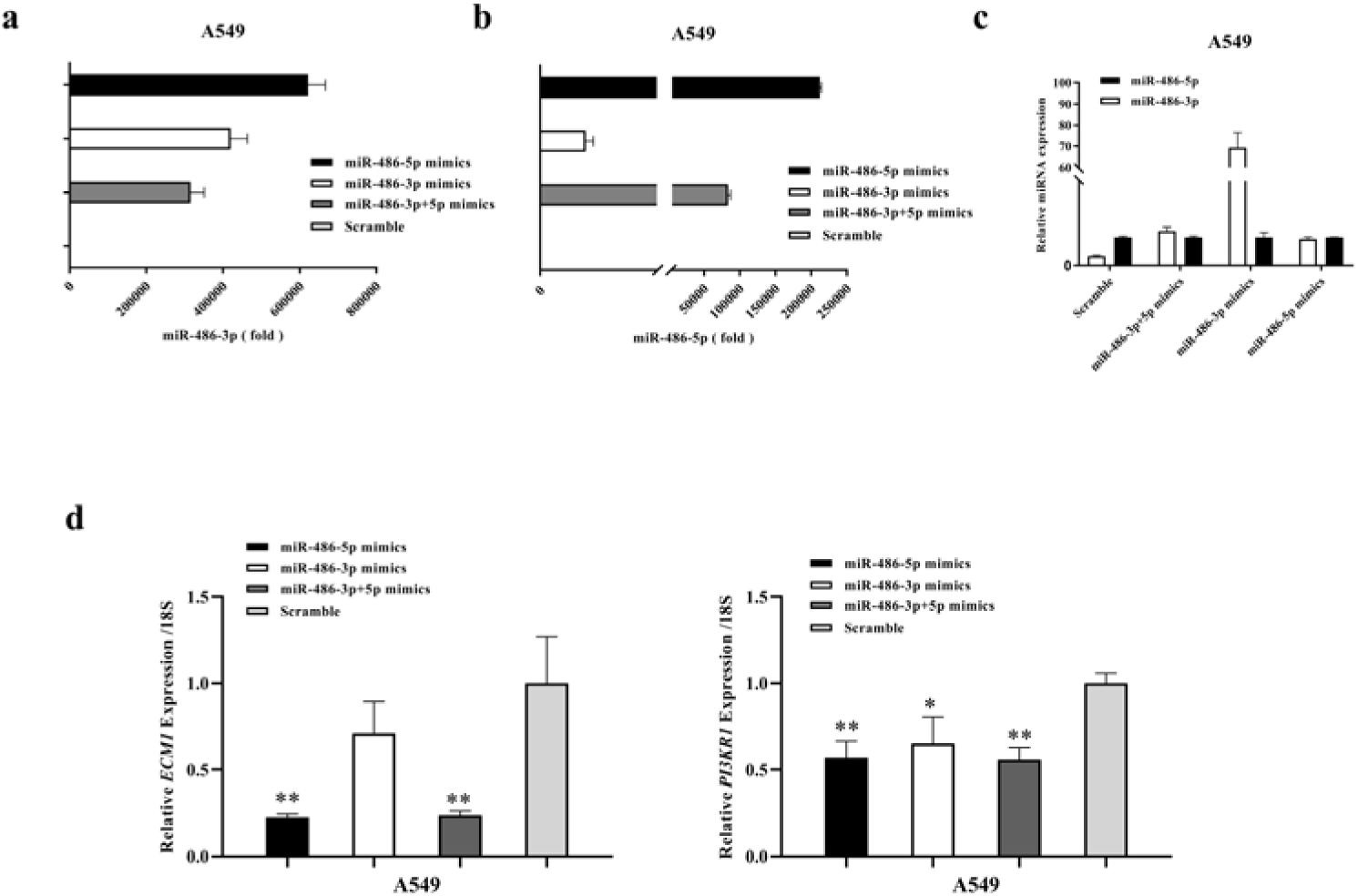
Simultaneous overexpression of miR-486-3p and miR-486-5p was detected, irrespective of whether miR-486-3p or miR-486-3p was used. The miR-486-3p mimics, miR-486-5p mimics, miR-486-3p and miR-5p mimic mix, and scramble control were transiently transfected into A549 cells. After 72 hours, the levels of mature miR-486-3p (a) and miR-486-5p (b) in RNA from A549 cells were determined by stem-loop quantitative real time RT-PCR. U6 snRNA served as an endogenous control. (c) The multiple, relative expression relationships of miR-486-3p and miR-486-5p were calculated, which used the expression of miR-486-5p as a reference. (d) The levels of the target gene (*ECM1, PI3KR1*) in the same RNA samples as (a, b) were determined by quantitative real-time RT-PCR. 18S RNA served as an endogenous control. \**P*<0.05 and \*\**P*<0.01 (Student’s *t*-test). The data are from three independent experiments (mean ± SEM).

### Lentivirus-mediated Pri-miRNA Expression for Specific miRNA Overexpression

Since the chemically produced miRNA–oligonucleotides are designed to be introduced into cells by transfection, the effects associated with their overexpression are invariably transient. We thus discovered that the cell viability changes with the influence of the transfection reagent’s toxicity. Therefore, the establishment of a method that would overexpress miRNA for a more prolonged period of time was thus very desirable and was needed to conduct a more detailed analysis of the networks formed between miRNA and the coding genes. In previous studies, researchers usually elucidated miRNA function by cloning the pri-miRNA sequences with 100–200 bp 5’ and 3’ flank sequences to the lentiviral vector with a cytomegalovirus (CMV) promoter (40,41). In order to verify the validity of this method for specific miRNA overexpression, we cloned the pri-miR-378 and pri-miR-486 sequences with 100 bp 5’ and 3’ flank sequences to the pLVSIN-CMV-Puro vector; the lentivirus that contains the pri-miR-378 and pri-miR-486 pairs was packed and the stable cell lines were constructed. We directly determined miRNA expression levels using stem-loop qRT-PCR. The pLVSIN-pri-miR-378 lentivirus (but not pLVSIN-pri-miR-486) induced the overexpression of miRNA (Figures 2). What is interesting is that miR-378-5p and miR-378-3p overexpression occurred simultaneously in the stable cell lines. Meanwhile, their target genes, *TOB2* and *Vimentin*, which were reported in previous studies(42,43), were downregulated respectively (Figure 2). As such, we can conclude that the method that cloned the pri-miRNA to the vector can be used to study the overall function of miRNAs, but it was not applicable to all miRNAs.

**Figure 2.**
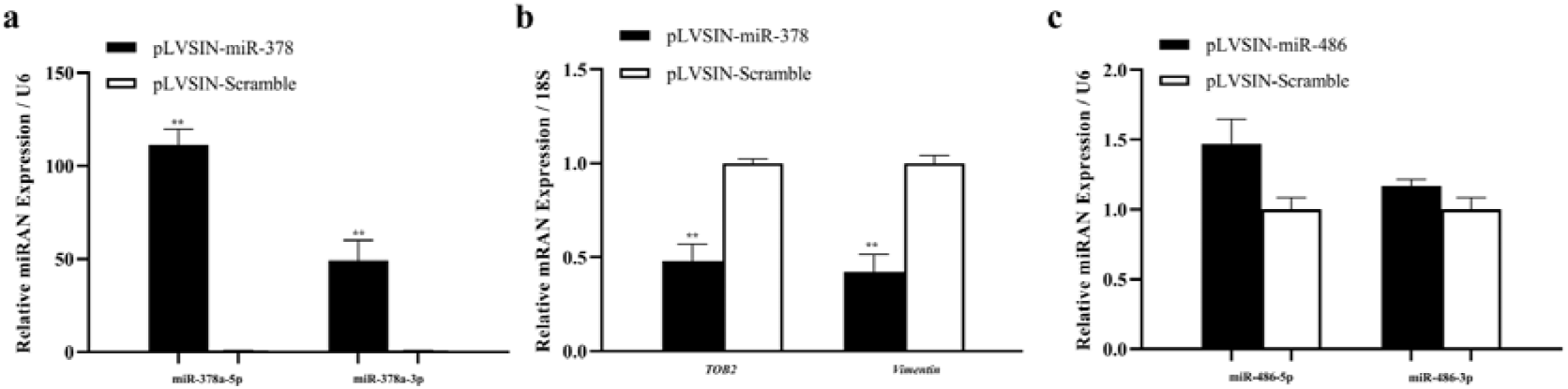
Specificity of the overexpressed effects of the pri-miRNA clone. pri-miR-378 and pri-miR-486 were cloned to the lentiviral pLVSIN CMV Puro vector, and the stably expressed pre-miR-378 and pre-miR-486 SKBR3 cell lines were constructed. (a) The expression of mature miR-378a-3p and miR-378-5p in the RNA of stably expressed pre-miR-378 SKBR3 cells was determined by stem-loop quantitative real-time RT-PCR. U6 snRNA served as an endogenous control. (b) The levels of the target gene (*TOB2* and *vimentin*) in the same RNA samples as (a) were determined by quantitative real time RT-PCR. 18S RNA served as an endogenous control. (c) The expression of mature miR-486a-5p and miR-486a-3p in RNA from stably expressed pre-miR-486 SKBR3 cells was determined by stem-loop quantitative real time RT-PCR. U6 snRNA served as an endogenous control. \**P*<0.05 and \*\**P*<0.01 (Student’s *t*-test). The data are from three independent experiments (mean ± SEM).

### Overexpression of Complementary miRNA Pairs Using Lentiviral Vectors Expressing Short-hairpin RNA

It is well-known that siRNA is a special form of miRNA that usually down-regulates target genes through a similar pathway. RNA interference (RNAi) technology using short-hairpin (sh)RNAs, which are transported to the cytoplasm by Exportin-5 and are processed by Dicer into siRNAs, has been detailed in past studies and has been widely exploited to modulate gene expression in a variety of mammalian cell types. Pre-miRNAs are structurally analogous to shRNAs and both are dependent on the karyopherin Exportin 5 (Exp5) for nuclear export. Pre-miRNAs reach the cytoplasm, at which point both the pre-miRNAs and shRNAs are processed by the RNase III enzyme Dicer to yield miRNA or siRNA duplex intermediates, one strand of which is then incorporated into RISC to downregulate the target gene [44, 45]. Considering the similarity of the processing mechanism for both pre-miRNA and shRNAs, we wonder whether the shRNA lentiviral platform can be used for specific miRNA overexpression. In our study, the pLKO.1-CTL vector, which is typically used for shRNA expression, was chosen to establish the lentiviral system to assess specific miRNA overexpression. Generally, when creating the shRNA cassette for a target gene, researchers usually suggest that the sense strand comes first, followed by the spacer, then the antisense strand [46]. We thereby designed a series of miRNA expression cassettes that placed the miR-486-3p sense downstream (#001) and upstream (#002) of the stem-loop.

We also constructed another miRNA expression cassette (#003), whereby the miR-486-3p sense in the downstream and upstream antisense sequences were mutated with four bases, as well as the miR-125a-5p expression cassette (#004), where mature sequences serve as the sense strand and have mismatches with the 3p strand; the latter was used as a control (Figure 3a). As shown in Figure 3b, we can detect the overexpression of miR-486-3p sense and antisense, irrespective of which comes first. However, with the mutation of the antisense upstream, the expression of the sense downstream significantly decreased. In addition, the expression of the miR-486-3p sense was higher than that of the antisense when we ensured that the miR-486-3p mature sequence sense came first when designing the miRNA-expression cassettes. More interesting is that whether the sense or antisense came first when we designed the miR-486-3p expression cassettes, all inhibited the proliferation of A549 cells through cell-cycle arrest and by promoting apoptosis (Figures 3c and 3d). This shares a similar function to the miR-486-5p that was reported in previous studies [19], while the miRNA expression cassette #003 plays the opposite role (Figures 3c and 3d). Mounting evidence suggests that the activation of wild-type p53 can promote the expression of miR-486 [19, 47]; thus, the impact of the addition of drugs that can activate p53 and stimulate the proliferation of A549 via stable miRNA expression cassettes was examined. As shown in Figure 4a, the addition of doxorubicin or nutlin3 all promoted the ability of miRNA cassettes to inhibit the proliferation of A549 cells, except in the case of miRNA expression cassette #003 which make the sensitivity of A549 cells to the anticancer drugs decreased. We wondered whether the opposite effect occurs due to the expression of the sense of miR-486-3p downstream of miRNA expression cassette #003 complementary to the mature miR-486-5p and then affect the function of target gene of miR-486-5p.

**Figure 3.**
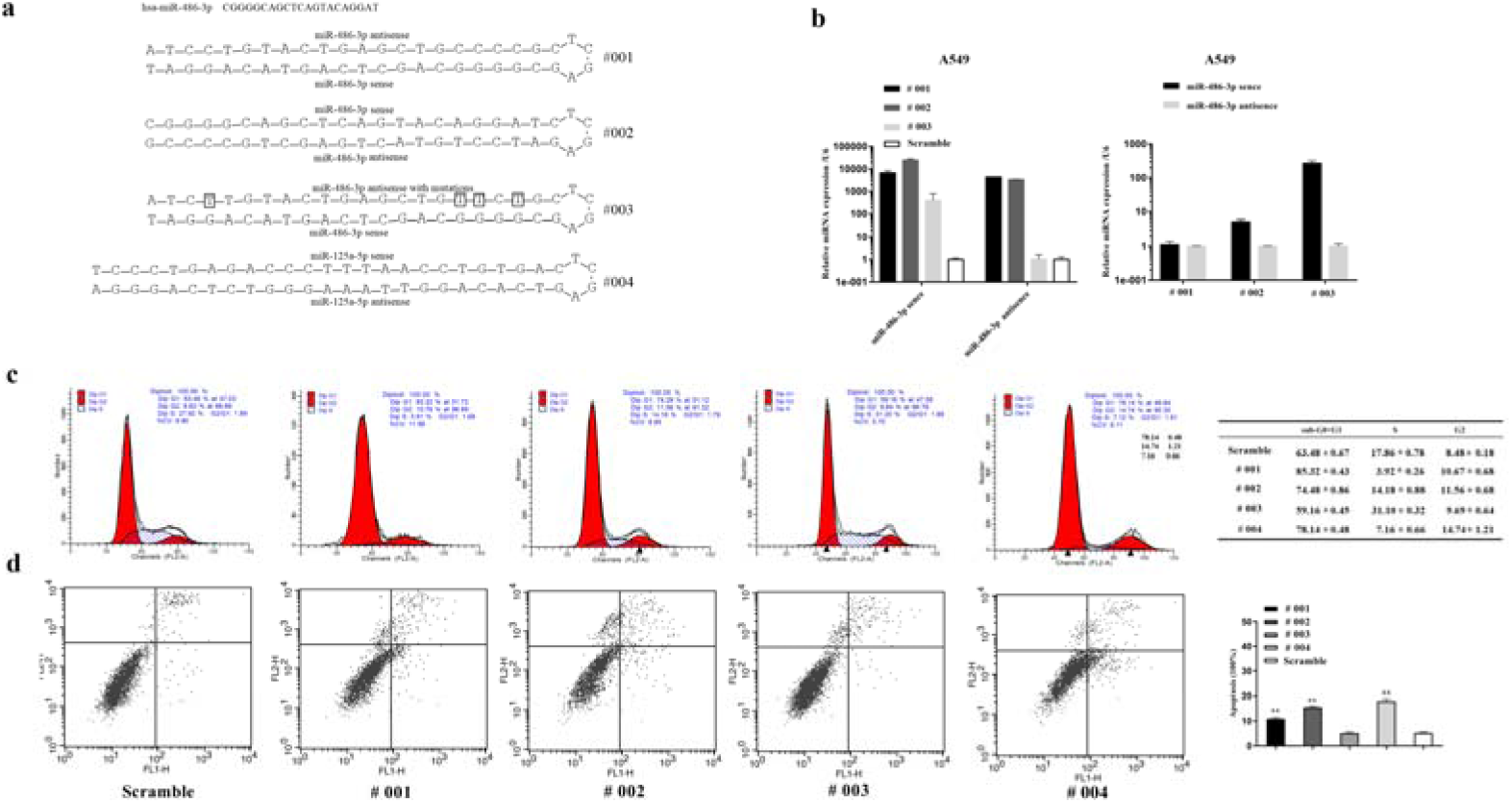
Specificity of the overexpressed effects of miRNA expression cassettes (#001–#004). The upstream and downstream strands of miRNA expression cassettes #001– #003 were composed by the miR-486-3p sense, antisense, and antisense with mutations while #004 consists of the miR-125a-5p sense and antisense. miRNA expression cassettes #001– #004 were cloned to the pLKO-TRC vector and the stable A549 cell lines were constructed. (a) The structure of miRNA expression cassettes #001–#004. (b) The level of sense and antisense in RNA from the stable A549 cell lines, which expressed the #001–#003 miRNA expression cassettes, was determined by stem-loop quantitative real time RT-PCR. U6 snRNA served as an endogenous control. (c, d) the effects of #001–#004 miRNA expression cassettes on the cell cycle and cell apoptosis in A549 cell lines. The cell line that stably expressed the scramble sequences were used as negative controls. \**P*<0.05 and \*\**P*<0.01 (Student’s *t*-test). The data are from three independent experiments (mean ± SEM).

**Figure 4.**
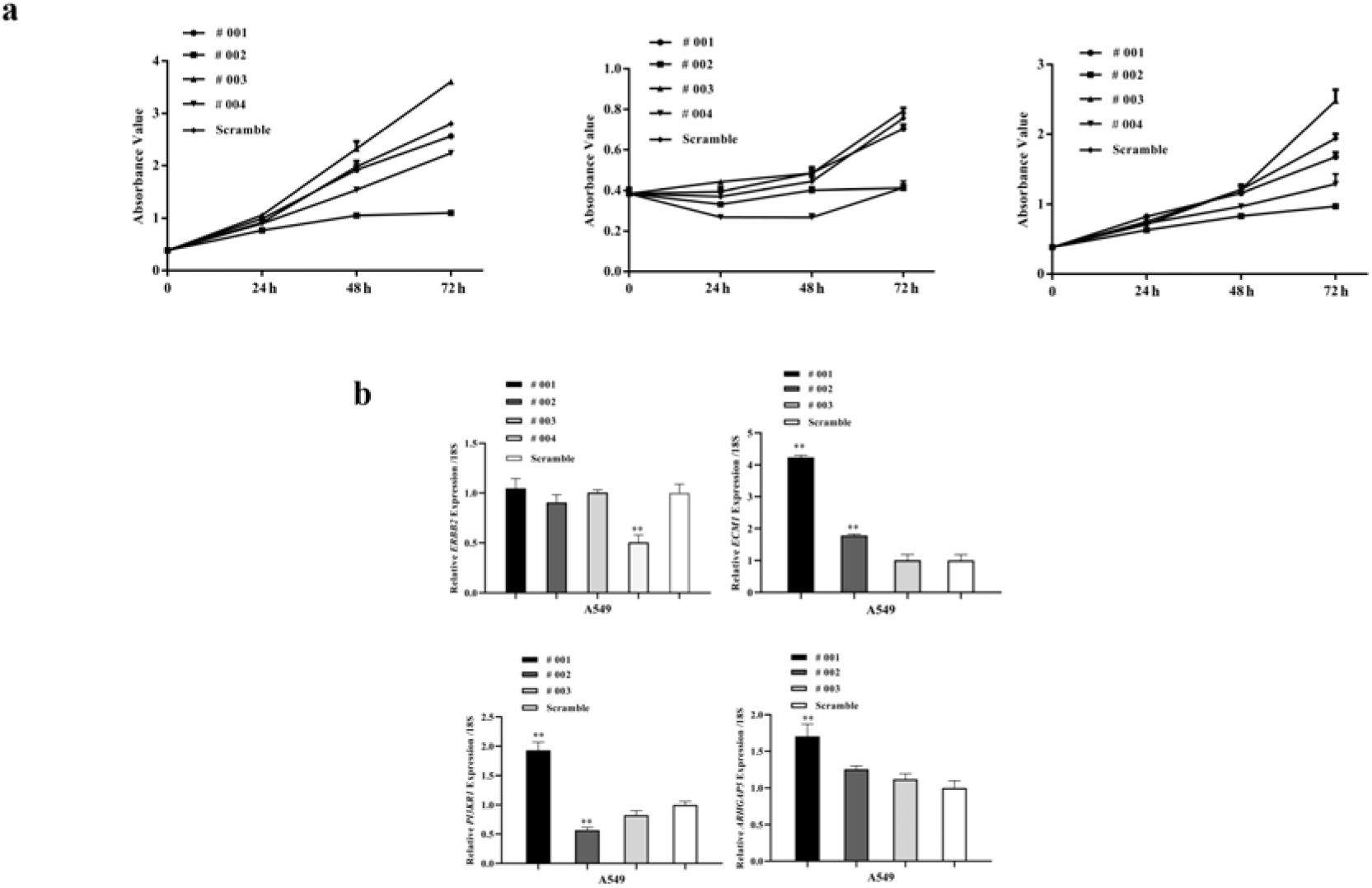
The effects of #001–#004 miRNA expression cassettes on cell proliferation and the target genes. (a) The effects of #001–#004 miRNA expression cassettes on cell proliferation, as related to the state of p53 activation, with the addition of doxorubicin and nutlin3. (b) Target gene (*Erbb2*, *ECM1*, *PI3KR1*, and *ARHGAP5*) expression, as reported by previous studies, was regulated by miR-125a-5p, miR-486-3p, and miR-486-5p, respectively, in stable A549 cell lines, which expressed the #001–#004 miRNA expression cassettes. The cell line that stably expressed the scramble sequences were used as negative controls. \**P*<0.05 and \*\**P*<0.01 (Student’s *t*-test). The data are from three independent experiments (mean ± SEM).

Thus, we then decided to detect the influence of miRNA expression cassettes (#001-#004) on the target genes *PIK3R1*, *ARHGAP5*, and *ECM1*, which are known to be regulated by miR-486-5p and miR-486-3p. However, the results did not match our expectations, as the *ECM1* and *PIK3R1* (which are reportedly downregulated by miR-486-3p and miR-486-5p) were significantly upregulated by the miRNA expression cassette #001. As such, the cassette #002 induced the higher expression of miR-486-3p sense when compared with the miR-486-3p antisense, and this downregulated the expression of *PIK3R1* but upregulated the expression of *ECM1* (Figure 4b). Further, the miR-125a-5p expression cassette, which has mismatches between the 5p and 3p strands, significantly reduced the expression of *Erbb2*, which is in agreement with the findings of previous studies [48]. To verify the results reported above, we also repeated the test on the H460 and HeLa cells (Figure S3). In addition, we also designed miRNA expression cassettes (#005 and #006) according to the miR-486-5p mature sequence; by doing so, similar results were obtained, irrespective of whether other cell lines were investigated or if the miR-486-5p expression cassettes were used (Figure 5 and Figure S4). From these results, we tentatively concluded that the miRNA expression cassettes, which were designed as miR-486-3p and miR-486-5p mature sequences, could be processed by the RNase III enzyme Dicer to express sense or antisense, and the expression levels of sense and antisense were related to the sequence. However, the results are inconsistent when the miRNA that we wish to overexpress possess complementary miRNA pairs, while the other miRNAs (which have mismatches between the 5p and 3p strands) are overexpressed correctly. As mentioned above, when creating the shRNA cassette for the target gene, researchers usually ensure that the sense strand comes first, followed by the spacer, and then the antisense strand. We hypothesize that the overexpression of sense upstream in the miRNA expression cassettes may play a major role in target-gene downregulation, while antisense overexpression inhibits the roles of sense.

**Figure 5.**
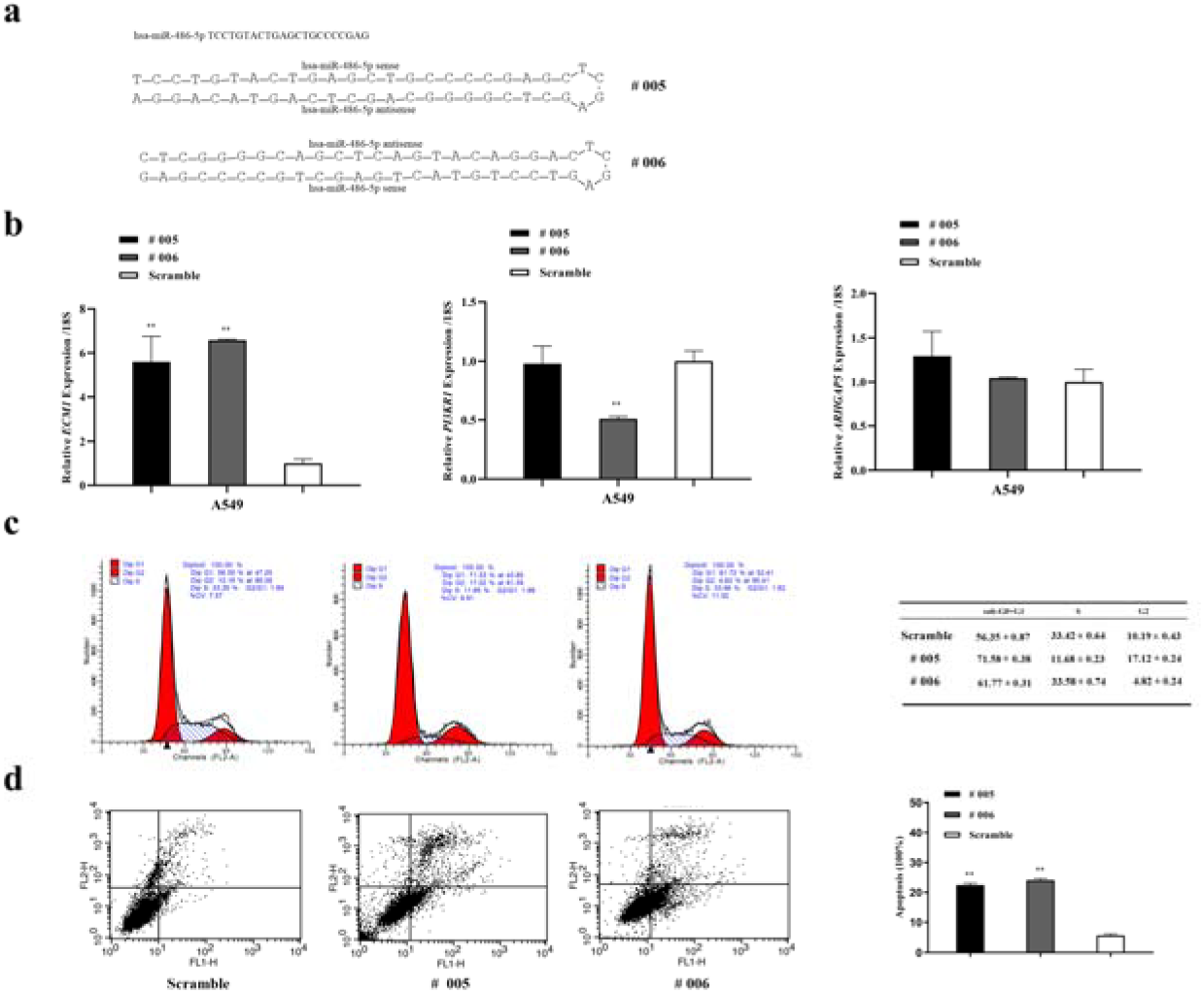
Specificity of the overexpressed effects of miRNA expression cassettes (#005–#006). The upstream and downstream strands of miRNA expression cassettes #005 and #006 were composed of the miR-486-5p sense or antisense. miRNA expression cassettes #005–#006 were cloned to the pLKO-TRC vector and the stable A549 cell lines were constructed. (a) The structure of the miRNA expression cassettes #005–#006. (b) The effects of #005–#006 miRNA expression cassettes on the expression of target genes *ECM1*, *PI3KR1*, and *ARHGAP5*. (c, d) The effects of #005–#006 miRNA expression cassettes on the cell cycle and cell apoptosis in A549 cell lines. The cell line that stably expressed the scramble sequences were used as negative controls. \**P*<0.05 and \*\**P*<0.01 (Student’s *t*-test). The data are from three independent experiments (mean ± SEM).

### Constructing Expression Cassettes for Specific miRNA Overexpression

Next, we designed a series of miRNA expression cassettes where the sense shared the same mature sequence of miR-486-3p (#007, #008, #009) and miR-486-5p (#010, #011), but the antisense was mutated with two or more loci, which was done in an attempt to eliminate the influence of the antisense in order to achieve the long-term overexpression of specific miRNAs (Figures 6a and 7a). The lentivirus, which contains the miRNA expression cassettes (#007, #008, #009, #010, #011), were packed and the A549 or H460 cells were infected to obtain stable cell lines. Then, we directly determined the miR-486-3p and miR-486-5p levels using quantitative real time RT-PCR. MiR-486-3p was overexpressed in the stable A549 cell line, which expressed miRNA expression cassettes #007, #008, and #009. While increasing the number of mutated bases of the downstream antisense sequence, the fold overexpression decreased (Figure 6b). Another notable finding was the downregulation of endogenous miR-486-5p expression, along with the overexpression of miR-486-3p; this is compliant with a rule where the increased multiple overexpression of miR-486-3p is consistent with the reduced multiple expressions of miR-486-5p. We also detected the expression of the target genes *PIK3R1*, *ARHGAP5*, and *ECM1* in the stable A549 cell line, which expressed the miRNA expression cassettes #007, #008, and #009. As shown in Figure 6c, the expression of *ECM1* and *PIK3R1* was significantly downregulated with the expression of miRNA expression cassettes #007, #008, and #009, which were either upregulated or did not change with the expression of the miRNA expression cassettes #001 and #002 with no mutation in the antisense downstream. Only the expression of miRNA expression cassette #007 induced the downregulation of *ARHGAP5*, while the other miRNA expression cassettes could not. Furthermore, we found that the cell cycles were significantly suppressed with the downregulation of the target gene. miRNA cassettes #007, #008, and #009 expressed in the H460 cells also yielded similar results (Figure S5). For miRNA expression cassettes #010 and #011 (Figure 7a), the overexpression of miR-486-5p could be detected and the expression of miR-486-3p was significantly suppressed (Figure7b). Further, we found that the target gene that was regulated by miR-486-3p in our study was upregulated with the overexpression of miR-486-5p (Figure 7b). This finding corresponds to the previous result. Moreover, we found that the overexpression of miR-486-5p could inhibit the proliferation of A549 cells, at least in part, by regulating the cell cycle (Figure 7c). However, the exact mechanism underlying this finding is not yet clear and requires additional investigation.

**Figure 6.**
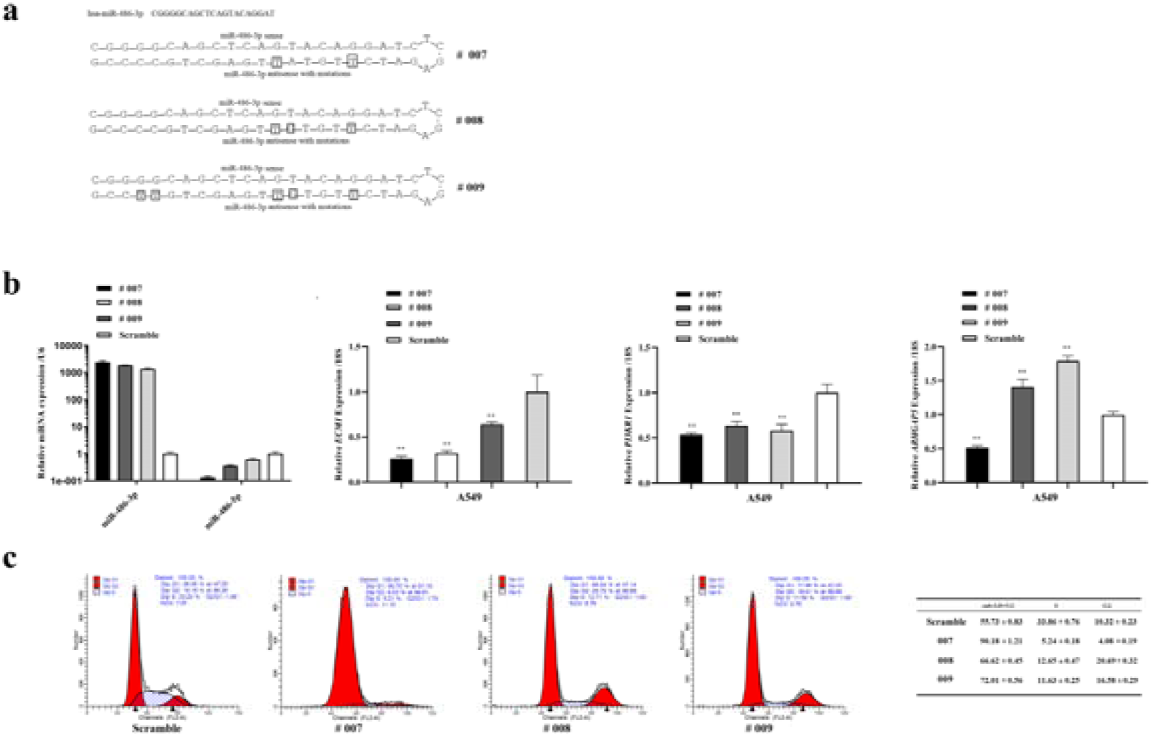
Specificity of the overexpressed effects of miRNA expression cassettes (#007–#009). The upstream and downstream strands of miRNA expression cassettes #007–#009 were composed of the miR-486-3p sense and antisense with mutations. The miRNA expression cassettes #007–#009 were cloned to the pLKO-TRC vector and the stable A549 cell lines were constructed. (a) The structure of miRNA expression cassettes #007–#009. (b) The expression of miR-486-3p and miR-486-5p and their target genes (*ECM1*, *PI3KR1*, *ARHGAP5*) in the stable A549 cell lines, which expressed the #007–#009 miRNA expression cassettes. (c) The effects of #007–#009 miRNA expression cassettes on the cell cycle of A549 cell lines. The cell line that stably expressed the scramble sequences were used as negative controls. \**P*<0.05 and \*\**P*<0.01 (Student’s *t*-test). The data are from three independent experiments (mean ± SEM).

**Figure 7.**
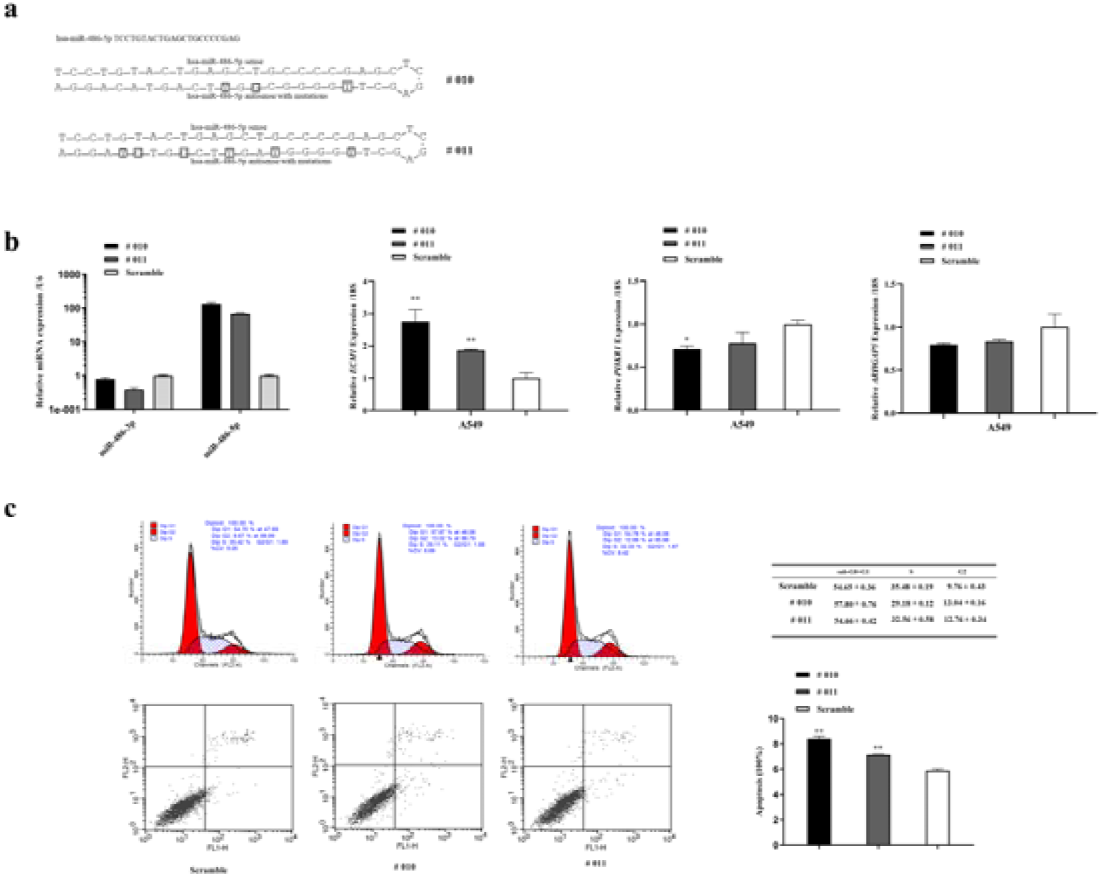
Specificity of the overexpressed effects of miRNA expression cassettes (#010–#011). The upstream and downstream strands of miRNA expression cassettes #010–#011 were composed of the miR-486-5p sense and antisense with mutations. The miRNA expression cassettes #010–#011 were cloned to the pLKO-TRC vector and the stable A549 cell lines were constructed. (a) The structure of miRNA expression cassettes #010–#011. (b) The expression of miR-486-3p, miR-486-5p, and their target genes (*ECM1*, *PI3KR1*, and *ARHGAP5*) in the stable A549 cell lines that expressed the #010–#011 miRNA expression cassettes. (c) The effects of #010–#011 miRNA expression cassettes on the cell cycle of A549 cell lines. The cell line that stably expressed the scramble sequences were used as negative controls. \**P*<0.05 and \*\**P*<0.01 (Student’s *t*-test). Data are from three independent experiments (mean ± SEM).

In order to further verify the miRNA expression cassettes function by expressing the upstream sense, rather than the mutation downstream from antisense, of the target gene. Expression cassettes #012, #013, #014, #015, and #016 were designed by mutating sense or antisense. As shown in Figure 8, we found the mutation of the miR-486-3p sense sequence; in this case, the miRNA expression cassettes still resumed their prominent roles in relation to the target gene. However, with the mutation of the miR-486-5p sense sequence, the miRNA expression cassettes lost their function to the target genes and cell cycle. From the results, we surmise that for some miRNAs, the mutation of the miRNA seed region sequence may have either a substantial, no, or little impact on the miRNA target function.

**Figure 8.**
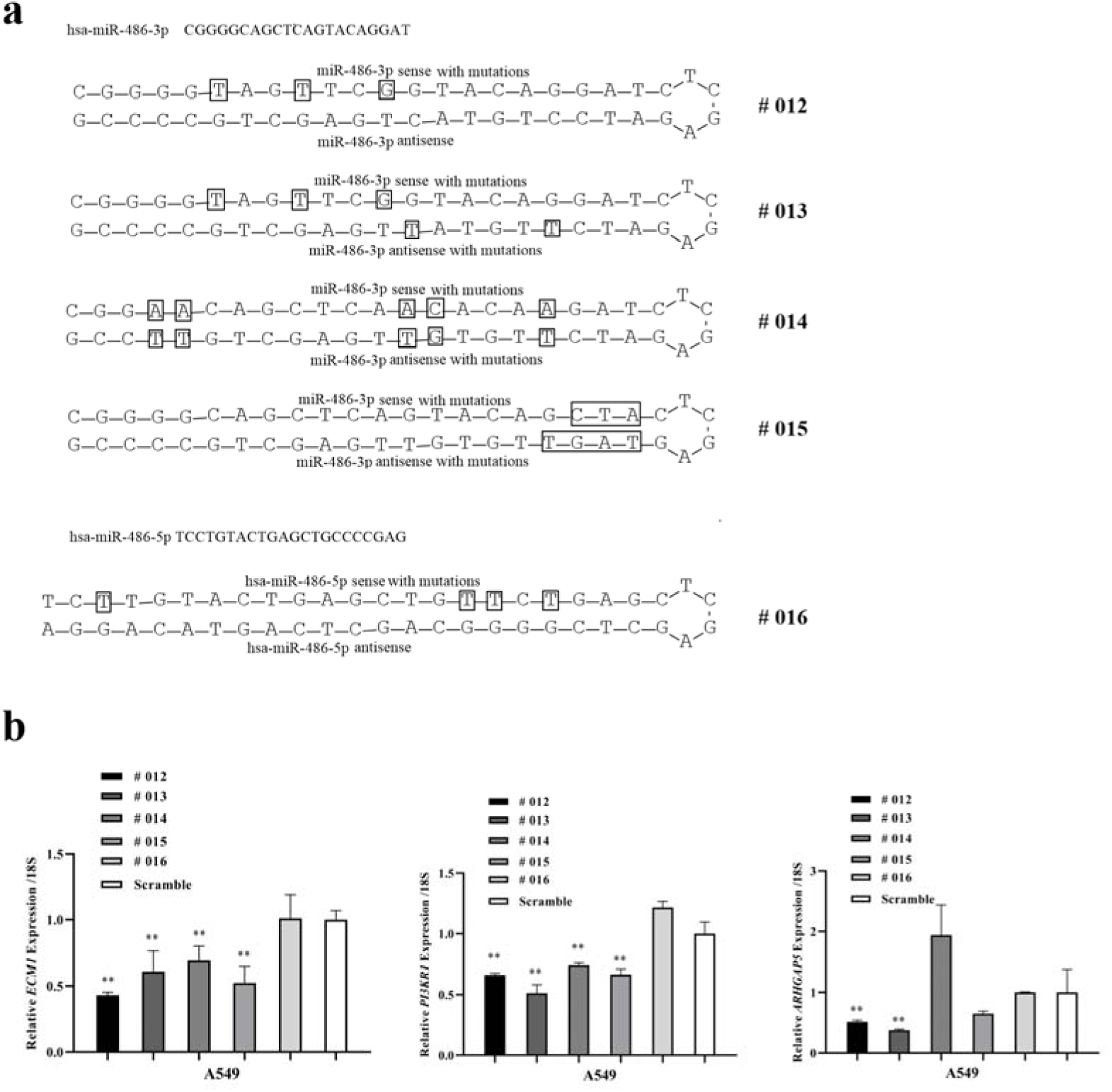
Specificity of the overexpressed effects of miRNA expression cassettes (#012–#016). The miRNA expression cassettes #012–#016 were cloned to the pLKO-TRC vector and the stable A549 cell lines were constructed. (a) The structure of the miRNA expression cassettes #012–#016. (b) The expression of the target genes (*ECM1*, *PI3KR1*, and *ARHGAP5*) in the stable A549 cell lines, which expressed the #012–#016 miRNA expression cassettes. The cell line that stably expressed the scramble sequences were used as negative controls. \**P*<0.05 and \*\**P*<0.01 (Student’s *t*-test). The data are from three independent experiments (mean ± SEM).

## Discussion

### The Discovery of Complementary miRNA Pairs had a Huge Impact on Traditional Methods

Researchers who have conducted previous studies in this field usually pay more attention to the specific miRNA activity being suppressed as the aberrant, high-level expression of certain miRNAs, which is frequently associated with the development of human cancers [10, 14, 16]. However, recent studies have shown that many miRNAs that function as potent tumor suppressors exhibit significant downregulation during cancer development [19, 22, 38]. It is indispensable to establish a method for the long-term overexpression of specific miRNA activity. Although chemically modified, double-stranded RNA miRNA mimics or angmir, which mimic endogenous miRNAs and pri-miRNA (usually expressed via RNA polymerase II or III) were used to overexpress miRNA in a study of their function [49–52]. Furthermore, studies of complementary miRNA pairs were also conducted. Based on our findings, we doubt the accuracy of traditional methods examining miRNA overexpression. Therefore, in our study, we chose miR-486-3p and miR-486-5p, which are complementary to each other, to verify the accuracy those methods. As shown in our study, miR-486-3p and miR-486-5p were significantly overexpressed in A549, H460, and HeLa cells, irrespective of whether the miR-486-5p or miR-486-3p mimics were transfected. These mimics were respectively overexpressed as well. Even more interesting is the fact that the target gene, which was downregulated by miR-486-5p, was also reduced by the transfection of miR-486-3p. We further discuss a method that used the pri-miRNA, which is expressed via RNA polymerase II. We found that the overexpression of miR-378-5p and miR-378-3p was detected, and that their associated target genes (as reported in previous studies) were also downregulated through the overexpression of pri-miR-378. Nevertheless, with respect to pri-miR-486, the overexpression of miR-486-5p was detected with no change in miR-486-3p. In all, the traditional methods adopted to overexpress miRNA is difficult to receive prospective specific miRNA activity increasing.

### Interaction between Complementary miRNA Pairs may Play New Regulatory Roles

RNAi technology has been studied in detail and widely exploited, and usually utilizes shRNA, which shares the same process mechanisms with pri-miRNA to modulate gene expression [10]. We designed many miRNA-expression cassettes in an attempt to achieve the correct approach to overexpress specific miRNA activity. The miRNA-expression cassettes usually take the mature sequence of miR-486-5p or miR-486-3p as the sense strand. When the antisense strand as their corresponding complementarity, the miRNA-expression cassettes all have influence on the cell cycle and cell apoptosis with no change or upregulate of the target gene of miR-486-3p and miR-486-5p reported by previous research and the expression of sense and antisense all can be detected. We are perplexed by these results, as there was an absence of target gene co-expression regulation of sense and antisense. We suppose that this function is the result of an interaction between sense and antisense; however, the detailed mechanism requires further exploration [30]. We also conclude that the overexpression of sense upstream in the miRNA expression cassettes may play a major role in target gene downregulation, while antisense overexpression inhibits the roles of sense, supporting the design principles of shRNA. This suggests that the sense strand comes first, followed by the spacer, and then the antisense strand [46].

### Complementary miRNA Pairs are Mutually Regulated

Furthermore, we also designed the miRNA expression cassettes, including antisense with mutations. With the antisense mutations being achieved according to the rules of T to C and A to G, the specific miRNA overexpression was testified in A549, H460, and HeLa cells. By adopting this approach, we suggest that according to the mutation rules (T to C and A to G), the bases of the antisense were mutated downstream, but with the presumption that the structure of the stem-loop was not broken down. With the destruction of the stem-loop, the sense and antisense might not be expressed and the shRNA or miRNA expression cassettes will lose their function. Moreover, the overexpression level of miRNA decreases to a greater degree with increases in the number of mutations in the antisense. The target gene of miR-486-5p is downregulated in cells that overexpress miR-486-3p. One point to note from our findings was that with the overexpression of miR-486-3p, the expression of miR-486-5p decreased. Conversely, increases in the multiple overexpression of miR-486-3p was also associated with the significant downregulation of miR-486-5p. This finding supports the results of a previous study, which proposed that the interaction between miRNA, its miRNA*, and other miRNA pairs with potential binding events may demonstrate evidence for a possible mutual regulatory pattern in a complex regulatory network [30, 31].

### The Mutation of the Seed Sequence of Certain miRNA May have Little Impact on its Function

Previous studies have shown that the region known as the seed sequence is considered to be a 6–8 nt long substring within the first 8 nt at the 5’-end of the miRNA [53, 54]. It is regarded as the most important feature for target recognition by miRNAs in mammalians [55, 56], as mutations in the seed regions of human miRNA usually lead to the development of many diseases by impacting miRNA biogenesis and resulting in a significant reduction of mRNA targeting [57, 58]. However, Recent studies also have revealed roles for miRNA sequences beyond this region in specifying target recognition and regulation [59]. In our study, we found that the effect of miR-486-3p never completely disappeared with mutations in the miRNA seed sequence, which is contrary to the expectation where the target gene would still be downregulated. While mutations of the miR-486-5p seed sequence, along with a loss of function, did not inhibit the proliferation and cell cycle of A549, particularly when two locus mutations were present in the seed sequence of miR-486-5p. Our results just support the two views at the same time. Given these properties, the methods used to achieve specific miRNA over-expression will be very useful for future studies in miRNA biology, as well as in the design of novel human gene therapies.

## Acknowledgements

This research was supported by the National Natural Science Foundation of China (Grant No.: 51773050) and the Natural Science Foundation of Heilongjiang Province of China (No. B2015005). English-language editing of this manuscript was provided by Journal Prep Services.

## References

[1] Bartel DP, MicroRNAs: genomics, biogenesis, mechanism, and function, CELL 2004; 116: 281–97;PMID:14744438;

[2] Carthew RW, Sontheimer EJ, Origins and Mechanisms of miRNAs and siRNAs, CELL 2009; 136: 642–55;PMID:19239886;10.1016/j.cell.2009.01.035

[3] Michlewski G, Caceres JF, Post-transcriptional control of miRNA biogenesis, RNA 2019; 25: 1–16;PMID:30333195;10.1261/rna.068692.118

[4] Gebert L, MacRae IJ, Regulation of microRNA function in animals, Nat Rev Mol Cell Biol 2019; 20: 21–37;PMID:30108335;10.1038/s41580-018-0045-7

[5] Wilczynska A, Bushell M, The complexity of miRNA-mediated repression, CELL DEATH DIFFER 2015; 22: 22–33;PMID:25190144;10.1038/cdd.2014.112

[6] Jonas S, Izaurralde E, Towards a molecular understanding of microRNA-mediated gene silencing, NAT REV GENET 2015; 16: 421–33;PMID:26077373;10.1038/nrg3965

[7] Rupaimoole R, Calin GA, Lopez-Berestein G, Sood AK, miRNA Deregulation in Cancer Cells and the Tumor Microenvironment, CANCER DISCOV 2016; 6: 235–46;PMID:26865249;10.1158/2159-8290.CD-15-0893

[8] Pinel K, Diver LA, White K, McDonald RA, Baker AH, Substantial Dysregulation of miRNA Passenger Strands Underlies the Vascular Response to Injury, CELLS-BASEL 2019; 8PMID:30678104;10.3390/cells8020083

[9] Flemming A, Heart Failure: Targeting miRNA pathology in heart disease, NAT REV DRUG DISCOV 2014; 13:336;PMID:24781545;10.1038/nrd4311

[10] Chakraborty C, Sharma AR, Sharma G, Doss C, Lee SS, Therapeutic miRNA and siRNA: Moving from Bench to Clinic as Next Generation Medicine, Mol Ther Nucleic Acids 2017; 8: 132–43;PMID:28918016;10.1016/j.omtn.2017.06.005

[11] Rupaimoole R, Slack FJ, MicroRNA therapeutics: towards a new era for the management of cancer and other diseases, NAT REV DRUG DISCOV 2017; 16: 203–22;PMID:28209991;10.1038/nrd.2016.246

[12] Ozcan G, Ozpolat B, Coleman RL, Sood AK, Lopez-Berestein G, Preclinical and clinical development of siRNA-based therapeutics, Adv Drug Deliv Rev 2015; 87: 108–19;PMID:25666164;10.1016/j.addr.2015.01.007

[13] Wittrup A, Lieberman J, Knocking down disease: a progress report on siRNA therapeutics, NAT REV GENET 2015; 16: 543–52;PMID:26281785;10.1038/nrg3978

[14] Haraguchi T, Ozaki Y, Iba H, Vectors expressing efficient RNA decoys achieve the long-term suppression of specific microRNA activity in mammalian cells, NUCLEIC ACIDS RES 2009; 37: e43;PMID:19223327;10.1093/nar/gkp040

[15] Scherr M, Venturini L, Battmer K, Schaller-Schoenitz M, Schaefer D, Dallmann I, et al., Lentivirus-mediated antagomir expression for specific inhibition of miRNA function, NUCLEIC ACIDS RES 2007; 35: e149;PMID:18025036;10.1093/nar/gkm971

[16] Gentner B, Schira G, Giustacchini A, Amendola M, Brown BD, Ponzoni M, et al., Stable knockdown of microRNA in vivo by lentiviral vectors, NAT METHODS 2009; 6: 63–6;PMID:19043411;10.1038/nmeth.1277

[17] Ebert MS, Sharp PA, MicroRNA sponges: progress and possibilities, RNA 2010; 16: 2043–50;PMID:20855538;10.1261/rna.2414110

[18] Cohen SM, Use of microRNA sponges to explore tissue-specific microRNA functions in vivo, NAT METHODS 2009; 6: 873–4;PMID:19935840;10.1038/nmeth1209-873

[19] Peng Y, Dai Y, Hitchcock C, Yang X, Kassis ES, Liu L, et al., Insulin growth factor signaling is regulated by microRNA-486, an underexpressed microRNA in lung cancer, Proc Natl Acad Sci U S A 2013; 110: 15043–8;PMID:23980150;10.1073/pnas.1307107110

[20] Zhou Q, Haupt S, Kreuzer JT, Hammitzsch A, Proft F, Neumann C, et al., Decreased expression of miR-146a and miR-155 contributes to an abnormal Treg phenotype in patients with rheumatoid arthritis, ANN RHEUM DIS 2015; 74: 1265–74;PMID:24562503;10.1136/annrheumdis-2013-204377

[21] Bhatia V, Yadav A, Tiwari R, Nigam S, Goel S, Carskadon S, et al., Epigenetic Silencing of miRNA-338-5p and miRNA-421 Drives SPINK1-Positive Prostate Cancer, CLIN CANCER RES 2018PMID:30587549;10.1158/1078-0432.CCR-18-3230

[22] Croset M, Pantano F, Kan C, Bonnelye E, Descotes F, Alix-Panabieres C, et al., miRNA-30 Family Members Inhibit Breast Cancer Invasion, Osteomimicry, and Bone Destruction by Directly Targeting Multiple Bone Metastasis-Associated Genes, CANCER RES 2018; 78: 5259–73;PMID:30042152;10.1158/0008-5472.CAN-17-3058

[23] Awan HM, Shah A, Rashid F, Wei S, Chen L, Shan G, Comparing two approaches of miR-34a target identification, biotinylated-miRNA pulldown vs miRNA overexpression, RNA BIOL 2018; 15: 55–61;PMID:29028450;10.1080/15476286.2017.1391441

[24] Weis BL, Guth N, Fischer S, Wissing S, Fradin S, Holzmann KH, et al., Stable miRNA overexpression in human CAP cells: Engineering alternative production systems for advanced manufacturing of biologics using miR-136 and miR-3074, BIOTECHNOL BIOENG 2018; 115: 2027–38;PMID:29665036;10.1002/bit.26715

[25] Lim MY, Ng AW, Chou Y, Lim TP, Simcox A, Tucker-Kellogg G, et al., The Drosophila Dicer-1 Partner Loquacious Enhances miRNA Processing from Hairpins with Unstable Structures at the Dicing Site, CELL REP 2016; 15: 1795–808;PMID:27184838;10.1016/j.celrep.2016.04.059

[26] Lal A, Navarro F, Maher CA, Maliszewski LE, Yan N, O’Day E, et al., qmiR-24 Inhibits cell proliferation by targeting E2F2, MYC, and other cell-cycle genes via binding to “seedless” 3’UTR microRNA recognition elements, MOL CELL 2009; 35: 610–25;PMID:19748357;10.1016/j.molcel.2009.08.020

[27] Anokye-Danso F, Trivedi CM, Juhr D, Gupta M, Cui Z, Tian Y, et al., Highly efficient miRNA-mediated reprogramming of mouse and human somatic cells to pluripotency, CELL STEM CELL 2011; 8: 376–88;PMID:21474102;10.1016/j.stem.2011.03.001

[28] Rupaimoole R, Wu SY, Pradeep S, Ivan C, Pecot CV, Gharpure KM, et al., Hypoxia-mediated downregulation of miRNA biogenesis promotes tumour progression, NAT COMMUN 2014; 5: 5202;PMID:25351346;10.1038/ncomms6202

[29] Okamura K, Phillips MD, Tyler DM, Duan H, Chou YT, Lai EC, The regulatory activity of microRNA* species has substantial influence on microRNA and 3’ UTR evolution, NAT STRUCT MOL BIOL 2008; 15: 354–63;PMID:18376413;10.1038/nsmb.1409

[30] Guo L, Sun B, Wu Q, Yang S, Chen F, miRNA-miRNA interaction implicates for potential mutual regulatory pattern, GENE 2012; 511: 187–94;PMID:23031806;10.1016/j.gene.2012.09.066

[31] Lai EC, Wiel C, Rubin GM, Complementary miRNA pairs suggest a regulatory role for miRNA:miRNA duplexes, RNA 2004; 10: 171–5;PMID:14730015;

[32] Kozomara A, Griffiths-Jones S, miRBase: annotating high confidence microRNAs using deep sequencing data, NUCLEIC ACIDS RES 2014; 42: D68–73;PMID:24275495;10.1093/nar/gkt1181

[33] Nawrocki EP, Burge SW, Bateman A, Daub J, Eberhardt RY, Eddy SR, et al., Rfam 12.0: updates to the RNA families database, NUCLEIC ACIDS RES 2015; 43: D130–7;PMID:25392425;10.1093/nar/gku1063

[34] Chen L, Heikkinen L, Wang C, Yang Y, Sun H, Wong G, Trends in the development of miRNA bioinformatics tools, BRIEF BIOINFORM 2018PMID:29982332;10.1093/bib/bby054

[35] Chen L, Heikkinen L, Wang C, Yang Y, Knott KE, Wong G, miRToolsGallery: a tag-based and rankable microRNA bioinformatics resources database portal, Database (Oxford) 2018; 2018PMID:29688355;10.1093/database/bay004

[36] Wang Z, The guideline of the design and validation of MiRNA mimics, Methods Mol Biol 2011; 676: 211–23;PMID:20931400;10.1007/978-1-60761-863-8_15

[37] Younger ST, Corey DR, Transcriptional gene silencing in mammalian cells by miRNA mimics that target gene promoters, NUCLEIC ACIDS RES 2011; 39: 5682–91;PMID:21427083;10.1093/nar/gkr155

[38] Wang J, Tian X, Han R, Zhang X, Wang X, Shen H, et al., Downregulation of miR-486-5p contributes to tumor progression and metastasis by targeting protumorigenic ARHGAP5 in lung cancer, ONCOGENE 2014; 33: 1181–9;PMID:23474761;10.1038/onc.2013.42

[39] Ye H, Yu X, Xia J, Tang X, Tang L, Chen F, MiR-486-3p targeting ECM1 represses cell proliferation and metastasis in cervical cancer, BIOMED PHARMACOTHER 2016; 80: 109–14;PMID:27133046;10.1016/j.biopha.2016.02.019

[40] Liu YP, Vink MA, Westerink JT, Ramirez DAE, Konstantinova P, Ter Brake O, et al., Titers of lentiviral vectors encoding shRNAs and miRNAs are reduced by different mechanisms that require distinct repair strategies, RNA 2010; 16: 1328–39;PMID:20498457;10.1261/rna.1887910

[41] Scherr M, Venturini L, Eder M, Lentiviral vector-mediated expression of pre-miRNAs and antagomiRs, Methods Mol Biol 2010; 614: 175–85;PMID:20225044;10.1007/978-1-60761-533-0_12

[42] Deng Z, Du WW, Fang L, Shan SW, Qian J, Lin J, et al., The intermediate filament vimentin mediates microRNA miR-378 function in cellular self-renewal by regulating the expression of the Sox2 transcription factor, J BIOL CHEM 2013; 288: 319–31;PMID:23135265;10.1074/jbc.M112.418830

[43] Feng M, Li Z, Aau M, Wong CH, Yang X, Yu Q, Myc/miR-378/TOB2/cyclin D1 functional module regulates oncogenic transformation, ONCOGENE 2011; 30: 2242–51;PMID:21242960;10.1038/onc.2010.602

[44] Carthew RW, Sontheimer EJ, Origins and Mechanisms of miRNAs and siRNAs, CELL 2009; 136: 642–55;PMID:19239886;10.1016/j.cell.2009.01.035

[45] Doench JG, Petersen CP, Sharp PA, siRNAs can function as miRNAs, Genes Dev 2003; 17: 438–42;PMID:12600936;10.1101/gad.1064703

[46] Keefe EPO, siRNAs and shRNAs: Tools for Protein Knockdown by Gene Silencing, Word Lab 2013

[47] Hall AE, Lu WT, Godfrey JD, Antonov AV, Paicu C, Moxon S, et al., The cytoskeleton adaptor protein ankyrin-1 is upregulated by p53 following DNA damage and alters cell migration, CELL DEATH DIS 2016; 7: e2184;PMID:27054339;10.1038/cddis.2016.91

[48] Scott GK, Goga A, Bhaumik D, Berger CE, Sullivan CS, Benz CC, Coordinate suppression of ERBB2 and ERBB3 by enforced expression of micro-RNA miR-125a or miR-125b, J BIOL CHEM 2007; 282: 1479–86;PMID:17110380;10.1074/jbc.M609383200

[49] Younger ST, Corey DR, Transcriptional gene silencing in mammalian cells by miRNA mimics that target gene promoters, NUCLEIC ACIDS RES 2011; 39: 5682–91;PMID:21427083;10.1093/nar/gkr155

[50] Trang P, Wiggins JF, Daige CL, Cho C, Omotola M, Brown D, et al., Systemic delivery of tumor suppressor microRNA mimics using a neutral lipid emulsion inhibits lung tumors in mice, MOL THER 2011; 19: 1116–22;PMID:21427705;10.1038/mt.2011.48

[51] Scherr M, Venturini L, Eder M, Lentiviral vector-mediated expression of pre-miRNAs and antagomiRs, Methods Mol Biol 2010; 614: 175–85;PMID:20225044;10.1007/978-1-60761-533-0_12

[52] Yue J, miRNA and vascular cell movement, Adv Drug Deliv Rev 2011; 63: 616–22;PMID:21241758;10.1016/j.addr.2011.01.001

[53] Liu H, Zhou S, Guan J, Identifying mammalian MicroRNA targets based on supervised distance metric learning, IEEE J Biomed Health Inform 2013; 17: 427–35;PMID:23192603;10.1109/TITB.2012.2229286

[54] Lewis BP, Shih IH, Jones-Rhoades MW, Bartel DP, Burge CB, Prediction of mammalian microRNA targets, CELL 2003; 115: 787–98;PMID:14697198;

[55] Bartel DP, MicroRNAs: target recognition and regulatory functions, CELL 2009; 136: 215–33;PMID:19167326;10.1016/j.cell.2009.01.002

[56] Nielsen CB, Shomron N, Sandberg R, Hornstein E, Kitzman J, Burge CB, Determinants of targeting by endogenous and exogenous microRNAs and siRNAs, RNA 2007; 13: 1894–910;PMID:17872505;10.1261/rna.768207

[57] Mencia A, Modamio-Hoybjor S, Redshaw N, Morin M, Mayo-Merino F, Olavarrieta L, et al., Mutations in the seed region of human miR-96 are responsible for nonsyndromic progressive hearing loss, NAT GENET 2009; 41: 609–13;PMID:19363479;10.1038/ng.355

[58] Hughes AE, Bradley DT, Campbell M, Lechner J, Dash DP, Simpson DA, et al., Mutation altering the miR-184 seed region causes familial keratoconus with cataract, AM J HUM GENET 2011; 89: 628–33;PMID:21996275;10.1016/j.ajhg.2011.09.014

[59] Chipman LB, Pasquinelli AE, miRNA Targeting: Growing beyond the Seed, TRENDS GENET 2019; 35: 215–22;PMID:30638669;10.1016/j.tig.2018.12.005

